# Klarigi: Characteristic Explanations for Semantic Data

**DOI:** 10.1101/2021.06.14.448423

**Authors:** Luke T. Slater, John A. Williams, Paul N. Schofield, Sophie Russell, Samantha C. Pendleton, Andreas Karwath, Hilary Fanning, Simon Ball, Robert Hoehndorf, Georgios V Gkoutos

## Abstract

**Background:** Annotation of biomedical entities with ontology classes provides for formal semantic analysis and mobilisation of background knowledge in determining their relationships. To date enrichment analysis has been routinely employed to identify classes that are over-represented in annotations across sets of groups, such as biosample gene expression profiles or patient phenotypes. These approaches, however, usually consider only univariate relationships, make limited use of the semantic features of ontologies, and provide limited information and evaluation of the explanatory power of both singular and grouped candidate classes. Moreover, they do not solve the problem of deriving cohesive, characteristic, and discriminatory sets of classes for entity groups.

**Results:** We have developed a new method, Klarigi, which introduces multiple scoring heuristics for identification of classes that are both compositional and discriminatory for groups of entities annotated with ontology classes. The tool includes a novel algorithm for derivation of multivariable semantic explanations for entity groups, makes use of semantic inference through live use of an ontology reasoner, and includes a classification method for identifying the discriminatory power of candidate sets. We describe the design and implementation of Klarigi, and evaluate its use in two test cases, comparing and contrasting methods and results with literature and enrichment analysis methods.

**Conclusions:** We demonstrate that Klarigi produces characteristic and discriminatory explanations for groups of biomedical entities in two settings. We also show that these explanations recapitulate and extend the knowledge held in existing biomedical databases and literature for several diseases. We conclude that Klarigi provides a distinct and valuable perspective on biomedical datasets when compared with traditional enrichment methods, and therefore constitutes a new method by which biomedical datasets can be explored, contributing to improved insight into semantic data.

## Background

The power of ontologies for representation of biomedical data has lead to ontological annotation becoming a preferred method for the characterisation of large biomedical datasets. Increasingly, domains such as medicine are adopting ontologies for annotation, aggregation, and analysis of patient data, opening up new methods for analysis hitherto unavailable using standard terminologies and nosologies. The ability to link data to these symbolic representations reduces inaccuracy and ambiguity, and permits integration with background knowledge through contextual aggregation. As such, biomedical ontologies and their instance data constitute a massive source of formalised knowledge.

This repository of formalised knowledge greatly facilitates secondary use, taking advantage of the semantic features of ontologies to integrate and explain the large and multi-modal datasets annotated to them. Semantic analysis methods are now applied to many knowledge synthesis and classification tasks, including prediction of protein-protein interactions and identification of rare disease genetic variants [15]. In the clinical space, semantic methods have been applied to tasks including diagnosis of rare and common diseases [26, 39] and identification of subtypes of diseases such as autism [44]. In addition, the synthesis of classical ontology-based methods and machine learning (ML) is seen as increasingly powerful [29].

Of great importance to the trust in, and consequently willingness to adopt, new analytical methods on the part of both the medical profession and the public, is the problem of explainability [21]. While application of an machine learning (ML) algorithm can generate a classifier based on large amounts of data, for example in patient population stratification, current methods make understanding the clinical basis for classification difficult. Despite the increasing use of semantics in biomedical analysis, the subsequent derivation of semantic explanations for classifications, outcomes, labels, or groups generated by those analyses, remains a challenging task, and a major practical, ethical, and technical issue [2, 6]. This is a problem for any new algorithmic approach, though current discussion has centred on the new ML approaches emerging in the health sciences.

We define the task of semantic explanation as the identification of characteristic sets of classes for groups of entities described by ontology classes. What constitutes a characteristic set of classes depends upon the parameters of a particular investigation, but can most often be defined by classes that are some combination of explanatory, compositional, and discriminatory.

One approach to semantic explanation is enrichment analysis. For example, and most prominently, gene set enrichment analysis (GSEA) identifies classes that are significantly over-expressed in a set of genes [10]. Enrichment analysis techniques have also been used to explore over-representation of terms in groups of entities associated with other datasets and ontologies [8, 48]. There are, however, multiple drawbacks to the use of enrichment analysis methods for semantic explanation:

- Traditional enrichment methods using a univariate statistical test provide limited information about the relationship of each explanatory term with the group of interest.
- They usually do not consider the multivariate contribution of sets of terms to a non-redundant characterisation of the group of interest.
- Most make little or no use of the semantic features of ontologies, or information-theoretic methods of evaluating candidate terms.
- They provide limited tools and metrics for evaluation of the overall explanatory power of groups of terms with respect to the group of interest.
- They can be difficult or impossible to use with non-preconfigured ontologies, often with a dedicated focus on the Gene Ontology (GO) [45].

While methods do exist to help correct for the dependence of statistical tests on related ontology classes, these post-hoc corrections do not take full advantage of the ontology to make inferential associations. Some methods such as XGR [9] and Func [35] do take ontology structure into account, with both requiring preprocessing beforehand if using ontologies other than GO (Func) or using ontologies outside a predefined set or without pre-computed annotation databases (XGR). In particular, OntoFunc provides inferential preprocessing for Func [16]. Recent Bayesian approaches to ontology enrichment likewise make use of ontology structure, but require additional numerical data to construct informative priors [18], or are restricted to the Gene Ontology [51]. Furthermore, most enrichment approaches consider only univariate relationships for individual terms; some investigations have approached multivariable enrichment, with one using a random walk approach on omics datasets [43], and another relying on pre-processing into principal components [47].

Moreover, enrichment methods are not intended to identify full and cohesive sets of characteristic terms for groups of annotated entities, and these methods therefore provide a limited solution to the task of semantic explanation. We propose to solve these problems by developing a new method for semantic explanation.

The method introduces metrics for quantification of multiple aspects of the relationship between classes, sets of terms, and groups of annotated entities. These include a configurable measure of information content, facilitating the involvement of information theoretic methods of evaluating candidates. This is combined with a new algorithm for derivation of multivariable explanatory sets of terms, optimised for discrimination and/or composition. The method makes use of ontological inference via an ontology reasoner, and can be used directly with any consistent OWL ontology without pre-processing. We also include native support for the Phenopacket format [19] to facilitate efficient use of Klarigi for large-scale import any analysis of clinical phenotype data using HPO.

In this work, we formally describe the method and its implementation. We discuss existing approaches and their potential limitations, explaining how our proposed approach solves these problems. We then describe development of the tool, Klarigi, that implements the method, resulting in an analysis framework that can be used to explore semantic datasets.

To evaluate our approach, we develop two use cases for characterisation of patient populations. We compare and contrast the results of the analysis with those of an enrichment analysis, and compare with existing literature and manually curated biomedical databases as gold standard. In doing this, we show that Klarigi can be used to provide insights into biomedical datasets via semantic explanation, providing a perspective that is both distinct and complementary to the use of enrichment analysis.

## Design and Implementation

The fundamental challenge of semantic explanation is to determine, given a set of groups, and sets of entities described by ontology classes associated with those groups, a set of characteristic classes for each group. Since we developed this approach to be applicable to any biomedical ontology, we will describe it, including a data model and algorithms, in abstract terms first. We will then describe their implementation into the Klarigi tool.

### Data Modelling

Klarigi aims to determine sets of classes that characterise a group of entities annotated by classes. Doing this requires, at minimum, a corpus with three features:

1. A set of entities
2. A set of groups, which are each associated with a set of entities
3. An OWL ontology describing all classes (terms) in the corpus

We can define these as sets, which will allow us to define metrics and heuristics on their contents. *G* is a set of groups, where *G_i_* is the set of entities ascribed to the group *i*. *E* is a set of entities, where *E_j_* is the set of classes associated with entity *j*. *O* is a set of classes in the ontology, where *O_x_* is a class in the ontology.

### Metrics

To identify ‘useful’ sets of explanatory classes for groups, we need to balance multiple qualities of the relationship between candidate classes and the group. Specifically, we want to identify classes that are, in comparison with other candidates:

1. Associated with greater proportion of entities in the considered group.
2. More associated with entities in the considered group than in other groups.
3. More informative and specific.

To measure these qualities and thus make them comparable, we define three univariate scores to measure them, encoding numeric measures of the explanatory power of each candidate class for the given group. We first define a simple function that enables us to determine whether or not a particular entity in the corpus is annotated with a class (or any of its subclasses):

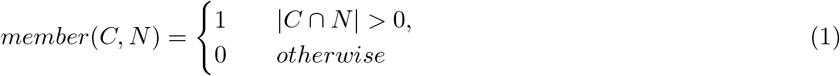

Given two sets *C* and *N*, any two sets of ontology classes, this function returns 1, if the size of the intersection of those sets is at least 1, and 0 otherwise. This allows us to identify whether any classes associated with a particular entity include our candidate class. Using this function, we can then define the first score, inclusion:

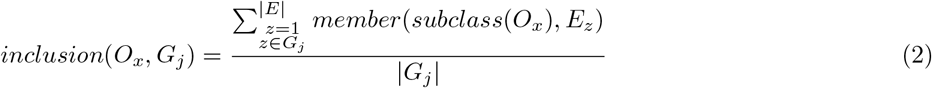

This score measures the proportion of entities in a given group that are annotated with either the candidate term or any of its subclasses. The *subclass* function is a call to the ontology reasoner, and returns a set of all transitive subclasses of the class passed as argument, including the original class. This enables the account of the deductive inferences made by the ontology reasoner, entailed by the axioms asserted in the considered ontology.

We next measure how uniquely a candidate class characterises the considered group, or its over-representation in the group with respect to others. Our previous work [40] defined a measure of exclusion (*exclusion_old_*) as:

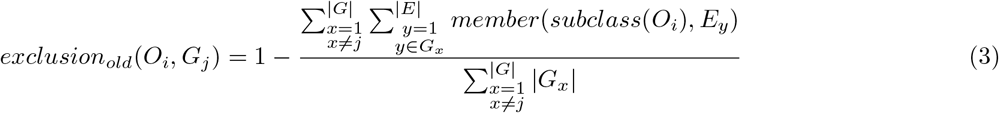

This provides a measure of the proportion of entities in the dataset that were associated with the class, but are attributed to *other* groups: the fewer entities external to the considered group that are annotated with the class, the greater the exclusion score. This measure, while useful in the context of previous experiments, it has certain drawbacks. In particular for the cases, where group sizes are unbalanced, the *exclusion_old_* measure may give rise to figures that are not easily comparable between different groups in the same dataset. For example, in the case of a dataset with 700 pneumonia patients and 300 pulmonary embolism patients, an exclusion score of 0.7 would be assigned with phenotype association with pneumonia, and 0.3 for pulmonary embolism, even if the inclusion was 0.5 for both of them; this is correct as a measure of absolute risk for the phenotype occurring in dataset, but is not necessarily helpful for exploring which phenotypes are more characteristic of the considered groups, in proportion to each other.

In GSEA, measurement of comparative expression is usually achieved by an *enrichment score,* which is usually defined by z-scores and/or odds and relative risk ratios, that measure the strength of association between the class and group membership; the ratio of B given the presence of A, and its symmetric inverse. However, their interpretation is challenging due to the behaviour of ratios and confusion between odds, relative, and absolute risk [7]. In the above example, the relative risk of both pneumonia and pulmonary embolism would be 1. However, this can become misleading in the presence of larger class imbalances [42]. For example, if we had 950 cases of pneumonia, and 50 cases of pulmonary embolism. If a phenotype appears thrice in this dataset, and two of those are associated with pulmonary embolism, we would get the extremely large values of 37.96 and 39.54 for relative risk and odds ratio respectively. While this is a valid association, it does not take into account the absolute risk of the event occurring with respect to the disparity in the group sizes, and so large values are given for uncommon and unlikely phenotypes, which can be easily misinterpreted. To address this issue, we define exclusion as:

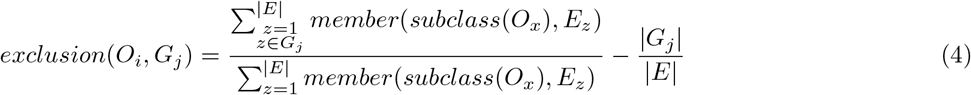

In this measure, the proportion of total entities in the corpus that are annotated with the considered class is determined. From this figure, the proportion of the total entities in the dataset that are associated with the current group is subtracted. This measures the overall representation of the class in the group versus the representation of the group in the overall population, and can be explained in terms intuitively similar to the calculation of an odds ratio: the odds of the event occurring in the exposed group minus the odds of exposure.

As such, it reflects the measurement of the representation of terms in a considered group, balanced with the absolute likelihood of the group appearing in the whole dataset. In the example above, the exclusion score for the phenotype with 0.5 inclusion with respect to both pneumonia and pulmonary embolism would be 0. This is because neither are more exclusively associated with the phenotype when their relative frequencies are taken into account.

In the latter example, our proposed phenotype association profile would lead to pulmonary embolism receiving an exclusion score of 0.617, and pneumonia an exclusion score of −0.61. These scores express the exclusion of the phenotype association with a group in the context of the overall appearance of the group in the dataset.

We further address the problem of balancing the internal and external characteristic power of candidate classes by introducing an additional heuristic to provide a balanced measure of our previously defined *inclusion* and *exclusion* scores. We note that the inclusion score is equivalent to the ‘absolute risk’ of the class being associated with an entity in the given group. On the basis of those two scores, we can further define:

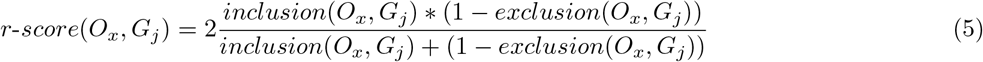

In the case of our latter example with a large class imbalance, our inclusion score for pulmonary embolism is 0.04, while it is 0.001 for pneumonia. Our *r-scores* would then be 0.036 and 0.0002, respectively. This provides a balanced metric between the internal representation of the class in the group, and its external exclusivity in the context of the larger dataset. Since both figures are taken into account, the r-score metric avoids extreme and potentially misleading values of z-score and odds ratio, when used to identify characteristic classes.

In the case where the program is being used with a corpus that only describes a single group, the *r-score* measure is alternatively defined as equivalent to *inclusion* (note that the program can be forced into this mode, see ‘exclusive group loading’ later). In this case, the algorithm does not consider exclusion at all, and only provides solutions in the context of *inclusion* – describing the internal composition of the group.

To test our third quality, we introduce a measurement of how informative or specific a candidate class is. Since this score does not rely on any relationship between groups and entities, it is calculated only once per ontology class:

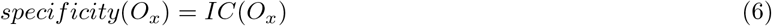

Specificity is defined as the result of a particular information content measure. These are configurable, and provided through implementation of the Semantic Measures Library [13], which implements many different information content measures. Currently, we implemented the Zhou [52] and Resnik [36] measures, which are defined as:

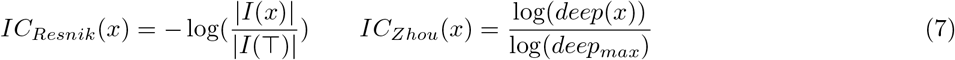

The Resnik measure is defined as the reverse log probability of the class appearing in the corpus, and will therefore be greater for classes that are annotated to entities more infrequently. The Zhou method is a measure of how deep the given class is when representing the ontology as a directed acyclic graph, calculated as a proportion of its maximum depth. Therefore, classes that are deeper in the ontology will have more intermediate subsuming classes between them and *Thing* and as a result will receive greater values. The choice of which information content measure to use is given to the user, and the best choice may differ depending on the dataset, ontology, or intended results [33].

### Scoring

Now that we have defined a number of measures for the explanatory power of candidate discriminating classes, the next steps depend upon identifying and scoring candidate classes in the relevant biomedical ontology. At this stage, we load and classify the ontology, verifying internal consistency and reflecting any structural inferences. Klarigi uses the ELK reasoner to perform this task [24], which supports the EL subset of OWL. Since we sought to support a maximal number of ontologies, datasets, and settings, we chose ELK since it ensures maximum polynomial time classification with respect to the number of axioms in the ontology. Since Klarigi is implemented using OWLAPI [17], however, use with more expressive reasoners can be easily implemented, or ontologies can be pre-processed before passing to Klarigi.

While scores can be derived, theoretically, for all classes in the biomedical ontology, in practice this is not necessary. This is because only classes that subsume classes appearing in our corpus can receive inclusion and exclusion scores above zero. We can therefore define a new set of candidate terms *C*, the set of candidate terms in the ontology that we will score, as a subset of *O*:

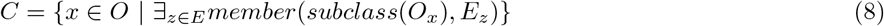

In practice, this is implemented more efficiently by iterating directly all classes appearing explicitly in the annotation corpus and identifying their transitive superclasses. Once *C* is determined, the scores are calculated as above for each combination of group and candidate class, excepting for specificity where the score is calculated only once per class.

### Candidate Restriction and Univariate Analysis

We have now created the set of *C* candidate explanatory classes, which consists of the set of all classes that either directly annotate, or subsume classes that directly annotate, all entities in the corpus. The next step is to identify characteristic candidate variables for each group of interest. We do this by creating a new set *U_j_* for each group:

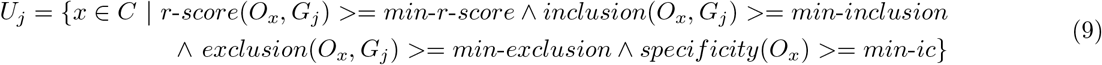

This new subset *U_j_* can be determined for each group, and contains a subset of the *C* candidate classes that meet the set of minimum restrictions for each score in relation to the considered group: r-score, inclusion, exclusion, and specificity. The process up to this stage consists the univariate mode of operation for Klarigi. At this point, the set *U_j_* can be output for a group of interest, and subsequently interpreted. The minimum cut-offs for scores are controlled by configurable parameters, described in Table 1.

**Table 1.** Names, descriptions, and default parameters for candidate class restrictions. These parameters define minimum values for the scoring heuristics, which restrict the set of candidate variables that will appear in the univariate analysis output, and be considered in the multivariable analysis stage.

| Parameter | Default | Description |
| --- | --- | --- |
| min-inclusion | 0 | Candidate terms with inclusion values below this level won't appear in, or contribute to, explanatory sets. |
| min-exclusion | 0.05 | Candidate terms with exclusion values below this level won't appear in, or contribute to, explanatory sets. |
| min-r-score | 0.1 | Candidate terms with *r-score* values below this level won't appear in, or contribute to, explanatory sets. |
| min-ic | 0.4 | Candidate terms with information content values below this level won't appear in, or contribute to, explanatory sets. |

### Multivariable Explanatory Sets

In addition to the univariate mode of operation, we also determine an additional method to identify sets of candidate terms that together characterise the group of interest. To do this, we can consider a solution space *S*, where *S_m_* is any potential subset of *U_j_*, the set of candidate classes that meet the minimum cut-offs for the univariate scores. Our goal is to identify such an *S_m_* that is a ‘good’ characterisation of entities in the group. To identify what constitutes a good characterisation of the group, we turn to our previously defined scores. For the purposes of our solution, we only consider *r-score* and *specificity* to evaluate the explanatory power of individual terms, relying on the ability of *r-score* to balance between *inclusion* and *exclusion* scores. To evaluate the overall fitness of a set of terms *S_m_*, we define an additional heuristic:

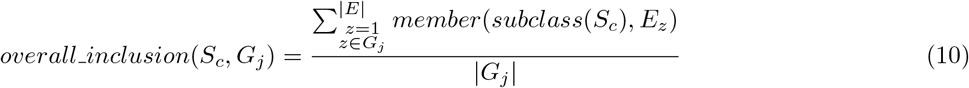

This is a measure of overall inclusion, the proportion of entities in the group of interest that are annotated by at least one of the classes in the proposed solution, or their subclasses. This measures how well a set of candidate classes covers the full set of entities in the group.

The challenge of obtaining a good solution lies with the optimisation of several variables, and can therefore can be considered as a multiple objective optimisation problem, considering the scoring heuristics as objective functions. The objective functions can be defined as the cut-offs for our scores. In the case of scores that measure the explanatory power of individual terms, this is the cut-off for candidate terms to appear in *S*, while the *overall_inclusion* function can be defined as the cut-off for *S* to be considered acceptable.

This can be considered in terms of the *ε*-constraints solution to multiple objective optimisation problems. In these solutions, one objective function is retained, while the rest are transformed into a set of constraints between which the remaining objective function can be optimised [12]. Our solution is inspired by this approach, selecting *overall_inclusion* as the objective function. However, instead of using static constraints for the other cutoffs, we develop an algorithm that steps down through acceptable values of these constraints in a priority order to dynamically identify high values for constraints in the context of objective function optimisation.

To do this, we define an order of priority for parameters:

1. *overall-inclusion*
2. *r-score*
3. *specificity*

The objective function can be considered as the highest priority constraint (and is also one that has no lower boundary). To identify a solution that maximises *overall_inclusion,* while optimising the other values within their configured boundaries, the algorithm steps down through acceptable values of each constraint in order of priority, from lowest to highest.

A satisfactory solution is defined as a set of ontology terms that meets a current minimum value for *overall_inclusion*, and in which every member meets the current minimum constraint for *r-score* and *specificity.*Upon each step of the algorithm, the subset of classes that meet the current individual criteria is identified, and if that set meets the current cutoff for *overall-inclusion,* this is returned as the solution. If not, the current constraint settings are stepped down in order of priority, with a new check for a satisfactory solution at each step. First, the cut-off for *r-score* is reduced stepwise to its lower limit, at which point the cut-off for *specificity* will be reduced by one step, and the cut-off for *r-score* will be reset. Once both *r-score* and *specificity* reach their lower limits, they are reset to their original values, and the *overall_inclusion* cut-off will be reduced by one step. This process is repeated until a satisfactory solution is found, checked every time a constraint is changed, or until the *overall_inclusion* cutoff reaches zero (a null response). The result of this process is a set of terms *S_good_*, a subset of *C*, that both maximises the value of the objective function, and maximises values of the constraints according to the order of priority. The algorithm is given in Algorithm 1.

The algorithm also supports maximal constraints, that are used to define the maximum score at which a candidate term can contribute to the *overall_inclusion* score. This is implemented in Algorithm 1 as the determination of the *S_pm_* subset of the proposed solution *S_p_*. By default, they are set to values that will not be triggered (shown in Table 2), but they can be configured to prevent the result set being dominated by a small number of highly explanatory classes, where a larger and more varied set is desired. For example, the appearance of the term `hypertension (HP:0008071)` at 0.95 inclusion for a group defined by a diagnosis of hypertension may not be very informative. Thus setting an upper limit to score values that can contribute to *overall_inclusion* will force the algorithm to continue to step down to find additional explanatory terms. Since these terms still characterise the group, however, they are still included in the explanatory set, and are therefore included in the output (and included in the final *overall_inclusion* value associated with the output). These upper parameters can be used to encourage the algorithm to seek larger, less monolithic explanatory sets.

**Table 2.** Descriptions and default values for multivariable parameters for the stepdown algorithm.

| Parameter | Default | Description |
| --- | --- | --- |
| max-exclusion | 1 | Candidate terms with r-score values above this level won't contribute to calculation of overall inclusion, though they will still appear in explanatory sets. |
| max-inclusion | 1 | Candidate terms with r-score values above this level won't contribute to calculation of overall inclusion, though they will still appear in explanatory sets. |
| max-r-score | 1 | Candidate terms with r-score values above this level won't contribute to calculation of overall inclusion, though they will still appear in explanatory sets. |
| top-ic | 0.7 | The maximum stepdown value for IC. |
| bot-ic | 0.4 | The minimum stepdown value for IC. |
| top-r-score | 0.8 | The maximum stepdown value for r-score. |
| bot-r-score | 0.1 | The minimum stepdown value for r-score. |
| top-total-inclusion | 1 | Maximum stepdown value for overall inclusion. |
| step | 0.05 | Cutoffs will be reduced by this step on each iteration of the stepdown algorithm. |

**Data:** Refer to Table 2.

**Result:** A set of ontology terms that explanatory of the cluster.

#### Algorithm 1: Algorithm for identifying characterising ontology terms for a cluster, by stepping down through r-score and specificity thresholds.

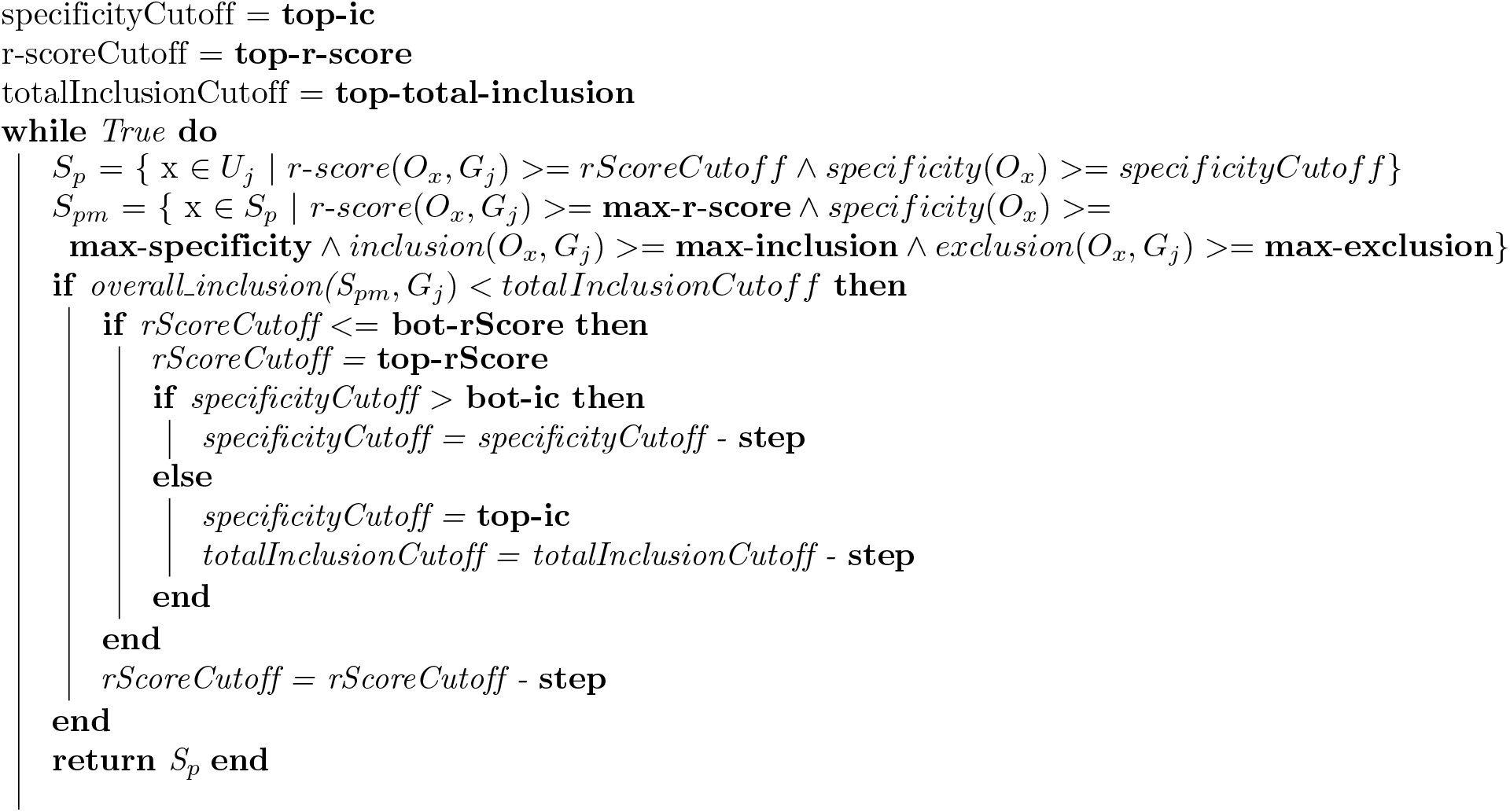

While the upper and lower boundaries for cut-offs are configurable, reasonable defaults have been set based on our observations using the algorithm. These can be defined for any of the term-wise scores: *r-score, inclusion, exclusion,* and *specificity.*

### Discrimination and Significance

On top of the univariate and multivariate modes of producing explanatory sets of classes for groups, we have also implemented several additional methods by which the fitness of solutions can be evaluated. To identify the discriminatory power of a set of explanatory classes, we can define a function that will, for each combination of entity and group in the corpus, produce a score:

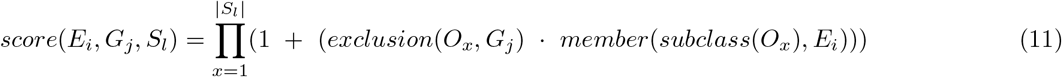

This score can either be calculated on the results of the multivariable stepdown approach, the full set of univariate results, or a manually specified list of term associations. In the case that we are using the univariate mode of execution only, *S_l_* will be equivalent to *U_j_*. Given a proposed solution for a group, consisting of a set of classes, a score is calculated for each entity in the corpus, defined as the product of one plus the exclusion score of every class in the solution set that the entity is annotated with.

This definition of a predictive score allows us to construct a model to classify group membership for entities, from which an Area Under the receiver operating Characteristic (AUC) score can be calculated. This provides a simple measure how well the set of explanatory variables, univariate or multivariate, was able to reclassify the set it was learned from. This can help to inform modifications to parameters or judge quality of the dataset in general.

The difference between these scores between the univariate and multivariate results can also provide insight on the quality of the modularisation performed by the stepdown algorithm. In the case that this value is small, there is a small amount of loss of total discrimination. Where values are large, there is a large decrease in performance when restricting scoring to the derived set of characteristic values, and this may be caused by a model that is poorly fit to the training data and does not reflect the full dataset.

The program also contains the possibility to apply this classifier to unseen data, calculating an AUC in the same way. This provides the ability to perform a test-set validation, or even external validation, to more properly judge the quality of the solution. Where this approach is applied to a new dataset, the *exclusion* score used is the one learned from the training corpus.

The second method of results evaluation is significance testing. In many instances, it is useful to access the type 1 error (false positive) rate to enable comparison with existing tools which provide p-values, and also to provide a conservative aid to interpreting inclusion and exclusion results. We implemented Monte Carlo methods to approximate the empirical p-value of the inclusion and exclusion statistics [34, 31]. Briefly, we sample with replacement a set of class associations for each entity of the same size as its original association set.

A test statistic (inclusion/exclusion) is generated for that set of patient profiles. This procedure is repeated 1,000 times to create a vector of test statistics. The statistics are ranked, and the empirical p-value is obtained as:

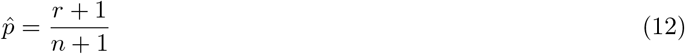

where *r* is the number of test statistics ranked greater or equal to the observed value, and *n* is the total number of permutations. The full algorithm is shown in Supplementary Algorithm 1.

The number of permutations can be changed by the user, and we suggest at least 1,000 to ensure a (log-)normal distribution and sufficiently precise p-values possible (minimum of 0.001 in this case). Throughout the evaluations in this article, we use 1,000 permutations. Since we are creating a set of p-values for non-independent phenotypes, we suggest a Bonferroni threshold be applied to determine an adjusted significance threshold:

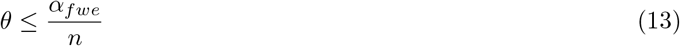

where *αfwe* is the desired error rate (0.05), *n* is the total number of phenotypes Klarigi generates p-values for per score. Thus *θ* is the new threshold for a significant inclusion/exclusion score.

### Output, Interpretation, and Configuration

Klarigi reports its results in either the univariate or multivariate mode, in the form of a table of class associations with each group of interest, reported with the scores assigned to them. These can either be received in a LATEXtable, a plain text table, or in Tab-Separated Values format. Each class is associated with four scores describing the relationship between the class and the group, which are given short descriptions in Table 3.

**Table 3.** Short description of Klarigi class scores and their interpretation.

| Score | Description |
| --- | --- |
| Inclusion | Measures the extent to which this term composes the group of interest. Greater values indicate a greater proportion of entities in the group being annotated with this class or its subclasses. |
| Exclusion | Measurement of how discriminative this term is for the group of interest. Greater values indicate that this class is more exclusively associated with the group. |
| r-score | A balanced measure of how compositional and discriminative the class is for the group. |
| IC | The information content of the class. The greater the value, the more informative or specific the class is, according to the chosen information content measure. |

The program is highly configurable, and we have previously discussed the parameters that are available to modify the operation and results of the program both for candidate restriction (see Table 1) and identification of multivariable modules (see Table 2). In addition, there are additional parameters that control different modes of operation and calculation of scores. For example, there is a choice between which information content measure to use for the calculation of the *specificity* score. Descriptions of parameters are available in the documentation for the program. Klarigi can also be run in exclusive group loading (EGL) mode, which only loads the group of interest, and does not consider any others. In this case, the stepdown algorithm operates upon *inclusion*, and neither *exclusion* nor *r-score* are calculated. This can be useful for investigating group constituency without taking into account the exclusivity of terms, and in the manuscript we subsequently refer to this as a compositional analysis, in contrast to the discriminatory analysis that results from loading all groups.

Because of this configurability, and the dependence of results on that configurability, we anticipate a general workflow of using the application to consist of interactive modification of those parameters according to intermediate results, and desired outcomes. For example, if an empty set is returned upon analysis, the **min-ic** parameter could be reduced in an attempt to identify explanatory terms with a lower *specificity* than previously considered.

## Results

We describe here the development and implementation of Klarigi, and make the tool freely available at https://github.com/reality/klarigi. This repository includes the source code, pre-compiled binaries, documentation, and a tutorial in notebook format that walks through the functionality of the software, explaining the significance of different parameters and outputs.

To demonstrate and evaluate the use of Klarigi for exploration of biomedical datasets, we developed two use-cases. Both describe clinical entities annotated using the Human Phenotype Ontology (HPO) [27]: in the first case text-derived phenotype profiles for a set of Medical Information Mart for Intensive Care III (MIMIC-III) admissions [22], and the second using a set of phenopackets [20], describing patients with rare diseases reported in literature [37]. We use Klarigi to explore both of these datasets, comparing and contrasting those results with the medical literature and enrichment analysis.

To further evaluate the results Klarigi produced for these use-cases, we also used an enrichment method to identify over-represented terms. To do this, we used XGR [9] to identify over-represented phenotypes using both one-tailed binomial and Fisher tests. P-values were adjusted with Bonferroni correction, propagating patient annotations to each term according to the True Path Rule. P-values were calculated with two background distributions: once using traits annotated to each patient and their frequencies as background, and additionally using only those annotated to the parent class of each entity tested; the maximum of each p-value was selected to help correct for the interdependence of testing. In the case of Klarigi, we also tested the normality of the permutation distribution for inclusion and exclusion scores by visually examining the distributions using histograms and QQ-plots. An R script containing these visualisations is provided in experiment repository.

The code used to obtain our results, as well as to perform the quantitative evaluations and create the tables for the paper, are available online at https://github.com/reality/klarigi_paper.

### Use Case: Pulmonary Embolism

Pulmonary embolism is a condition in which a thrombus (usually a blood clot) forms in, or migrates from a distal site, to the pulmonary arteries and occludes the pulmonary circulation, blocking the entry of blood into or through the lungs. This may be associated with predisposing conditions such as cancer, cardiovascular disease or lung disease, but is often sporadic, for example following a deep vein thrombosis in the leg. This condition is associated with a considerable mortality rate, and often presents acutely in the emergency room in ways that render it difficult to diagnose when associated with other co-morbidities, such as COPD, and typically shares symptoms with other more common conditions, such as pneumonia and acute bronchitis [32]. The time-critical dependence of treatment on diagnosis makes it important to identify combinations of discriminating symptoms as rapidly as possible [4].

To demonstrate Klarigi’s functionality, and to gain insight into the phenotypic presentations associated with pulmonary embolism and pneumonia, we created and evaluated text-derived phenotype profiles describing critical care admissions with the two diagnoses. We identified relevant admissions from MIMIC-III, and extracted phenotype profiles from their associated discharge letters. MIMIC-III is a large freely accessible dataset concerning nearly 60,000 visits to the emergency department at the Beth Israel Deaconess hospital [22].

We used the ICD-9 codes `486` and `41519` to identify 5,437 MIMIC-III admissions with either pneumonia or pulmonary embolism as an associated diagnosis (provided in the coded diagnosis list produced by a coding expert): 4,853 with pneumonia and 912 with pulmonary embolism. We further restricted these to admissions with pneumonia or pulmonary embolism given as the primary diagnosis, and those which did not have the other diagnosis in any position. This led to a final set of 991 admissions, made up of 699 primarily coded with pneumonia, and 292 with pulmonary embolism. We employed the 2021-08-02 release of the Human Phenotype Ontology, available at http://purl.obolibrary.org/obo/hp/releases/2021-08-02/hp.owl.

To create phenotype profiles for these admissions, we used the Komenti semantic text-mining framework [38], which implements Stanford CoreNLP [30]. For every considered admission, we collected the text from the corresponding discharge note. We created a lemmatised vocabulary of all labels and synonyms in the Human Phenotype Ontology (HPO) [27], containing 50,265 labels for 16,019 unique classes. We used these to identify ontology terms mentioned in the discharge notes. We then removed all associations that were identified by Komenti as negated or uncertain. We also removed classes equivalent to, or subclasses of, `pneumonia (HP:0002090), pulmonary embolism (HP:0002204)`, and `abnormal thrombosis (HP:0001977)`. To facilitate a holdout validation, we then reserved a randomly sampled 20% of the annotated admissions. In the training set there are a total of 799 records, 568 pneumonia and 232 pulmonary embolism, while in the test set there were 190 records, 60 pulmonary embolism and 131 pneumonia.

In its univariate mode, Klarigi calculates all relevant metrics for every class that contains an instance in the dataset, restricted by the options given in Table 1. The results of this analysis are shown in Supplementary Tables 1 and 2. The subsequent results form different modular subsets of these univariate results, by using the stepdown algorithm with reference to different objective functions, as well as potentially restricting the initial set of univariate results by modifying minimum scoring thresholds.

We then employed Klarigi to perform a compositional analysis of both groups separately, using “exclusive group loading” (EGL) mode to consider each group without reference to its relationship to any other groups in the dataset. As such, only the inclusion and IC scores are used, and the step-down algorithm uses inclusion rather than r-score as its optimisation parameter. The results of this analysis are shown in Table 4. For both conditions, large values of the overall inclusion score show that the given explanatory sets describe almost all of the group. Large values for *inclusion* on individual classes also indicate terms that cover a large proportion of the relevant group.

**Table 4.** Compositional analysis performed for pneumonia and pulmonary embolism using Klarigi. This is intended to identify a set of variables that characterise each group separately. This is achieved by using Klarigi in Exclusive Group Loading (EGL) mode. As such, the program produces a set of compositional classes for each disease, without considering how or whether those classes can be used to distinguish between the two groups. These results are a subset of the univariate results (visible in Supplementary Tables 1 and 2) sufficient to meet the threshold for *overall_inclusion.* Resnik IC was used. *max-inclusion* was set to 0.7 for pneumonia, to avoid returning only `dyspnea (HP:0002094)` and `abnormal breath sound (HP:0030829)`, which alone meet a sufficient *overall_inclusion. min-inclusion* was set to 0.3 for pulmonary embolism to reduce the size of the result set.

| pneumonia | Inclusion | IC | pulmonary embolism | Inclusion | IC |
| --- | --- | --- | --- | --- | --- |
| Dyspnea (HP:0002094) | 0.8 | 0.83 | Pain (HP:0012531) | 0.81 | 0.63 |
| Abnormal systemic blood pressure (HP:0030972) | 0.76 | 0.76 | Abnormal pattern of respiration (HP:0002793) | 0.77 | 0.7 |
| Abnormal breath sound (HP:0030829) | 0.69 | 0.8 | Abnormal systemic blood pressure (HP:0030972) | 0.76 | 0.76 |
| Cough (HP:0012735) | 0.67 | 0.87 | Dyspnea (HP:0002094) | 0.75 | 0.83 |
| Increased blood pressure (HP:0032263) | 0.64 | 0.83 | Increased blood pressure (HP:0032263) | 0.66 | 0.83 |
| Hypertension (HP:0000822) | 0.64 | 0.89 | Hypertension (HP:0000822) | 0.66 | 0.89 |
| Abnormality of temperature regulation (HP:0004370) | 0.48 | 0.77 | Abnormality of fluid regulation (HP:0011032) | 0.63 | 0.67 |
| Fever (HP:0001945) | 0.48 | 0.83 | Edema (HP:0000969) | 0.62 | 0.67 |
| *Overall* | *99.3* | - | Arrhythmia (HP:0011675) | 0.61 | 0.64 |
|  |  |  | *Overall* | *100.0* | - |

To gain insight on the differences between the phenotypic composition of the two groups, we also employed Klarigi to perform a discriminative analysis. In this case, it uses the opposing group as context, providing the additional *exclusion* and *r-score* measures, using the latter as the optimisation parameter for application of the stepdown algorithm. These results are shown in Table 5. We tested different hyper-parameter options based on the reclassification AUC on the training set, but finally used the default values, obtaining final values of 0.903 and 0.996 for pulmonary embolism and pneumonia respectively. Permutation testing was performed after hyper-parameter optimisation, and so was only run once. Meanwhile, the results of the enrichment analysis using both the Fisher and binomial tests are shown in Supplementary Table 3.

**Table 5.**
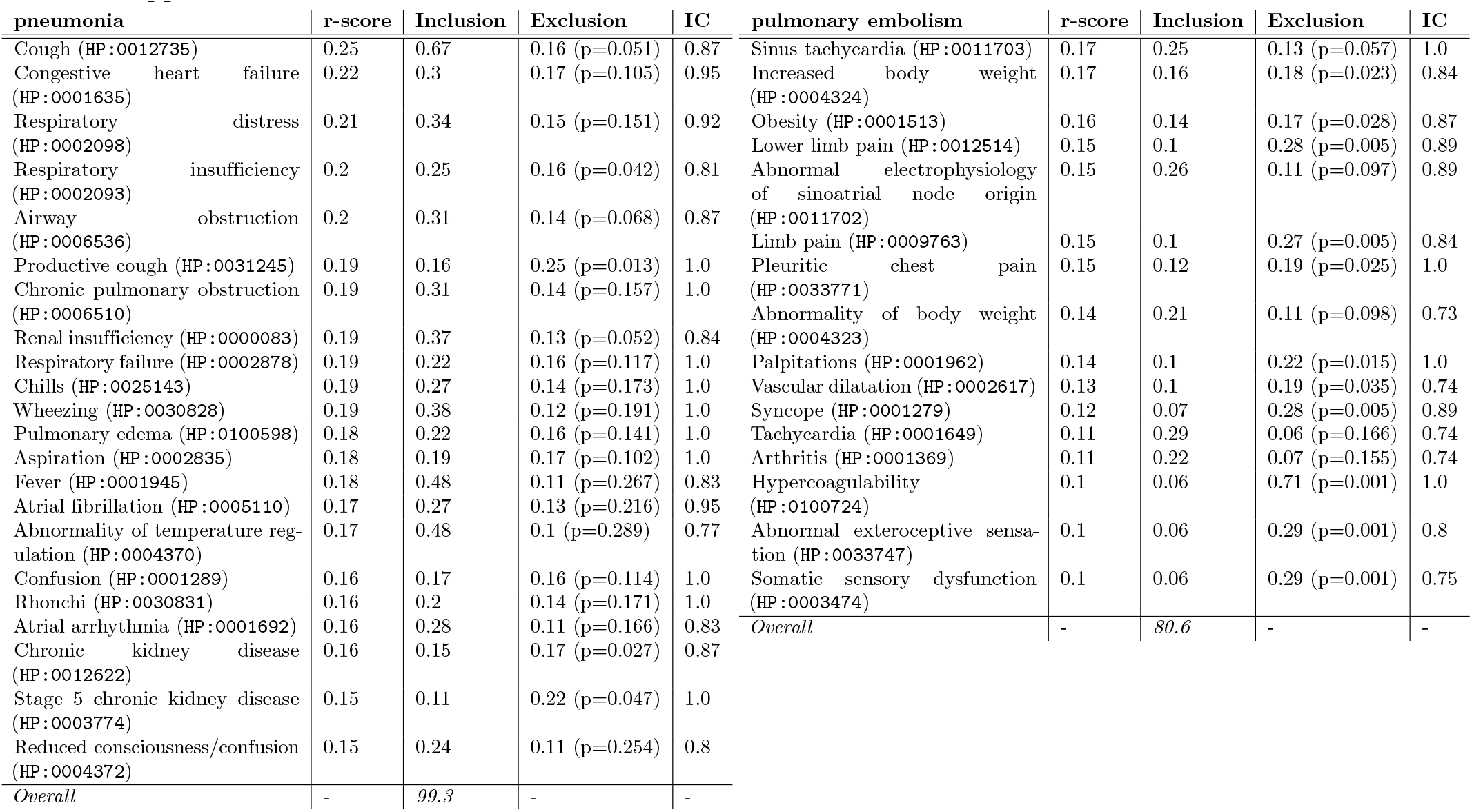
Results for the discriminatory analysis performed using Klarigi. Here, we are attempting to identify an exclusively compositional set of explanatory classes for each group, containing classes that can be used to discriminate between the two groups, introducing the *exclusion* and *r-score* measures to achieve this. These results are contrasted with the purely compositional analysis seen in Table 4, since it forms an alternative subset of the univariate results shown in Supplemental Tables 1 and 2.

Scores for *inclusion* measure the proportion of entities in the group - in this case PE or pneumonia - that are annotated by the class. Scores for *exclusion* indicate the extent to which an individual annotated to that class is likely to belong to one group rather than another, e.g. more likely to have pneumonia or PE, or vice versa. This is therefore a set of discriminating features that can be used to allocate a patient to one group or the other.

To evaluate how characteristic and discriminatory the explanatory sets were of their respective diseases, we used the 20% holdout set to define a classification task. The results of this analysis are listed in Table 6. In this analysis, we used the sets of characteristic terms identified by Klarigi in univariate mode, Klarigi in multivariate mode, and enrichment. Since the binomial enrichment results were a subset of the Fisher results, we combine these into one ‘enrichment’ set of explanatory terms. We then constructed classifiers for pulmonary embolism and pneumonia in the test set with scores for each entity identified using Klarigi. To provide additional measures of evaluation, as well as to evaluate the utility of the exclusion score in measuring term discrimination, we also compared the Klarigi classification method with semantic similarity approaches. To do this, we also built classifiers scored using the groupwise similarity between each phenotype profile in the test set, and the explanatory set for each method. We performed this task for both the best match average and average groupwise strategies, with both cases using the Resnik pairwise method and Resnik information content measure (using the annotations in the training set for the probability distribution). For all classifiers, we calculated an AUC using the scores for each admission in the test set, and their actual diagnosis group for the label.

**Table 6.** Test set classification results for pneumonia and pulmonary embolism, using Klarigi’s classifier (see Equation 11), and semantic similarity using the Resnik IC and Resnik pairwise method, with results given for both best match average (SS-BMA) and average groupwise (SS-AVG) strategies. OI is the overall inclusion score, defined in Equation 10, of the solution with respect to the test corpus. The greatest score for each category is emboldened.

| Group | Method | AUC (Klarigi) | AUC (SS-BMA) | AUC (SS-AVG) | OI |
| --- | --- | --- | --- | --- | --- |
| Pneumonia | Enrichment | 0.989 | 0.748 | 0.714 | 0.977 |
|  | Klarigi (u) | **0.996** | **0.769** | 0.73 | **0.992** |
|  | Klarigi (mv) | 0.992 | 0.771 | **0.752** | 0.985 |
| Pulmonary embolism | Enrichment | 0.575 | 0.543 | 0.561 | 0.15 |
|  | Klarigi (u) | **0.858** | **0.631** | **0.675** | **0.716** |
|  | Klarigi (mv) | 0.825 | 0.594 | 0.661 | 0.65 |

### Use Case: Rare Disease Annotation with Phenopackets

Phenopackets are a standardised format for the representation of phenotypic descriptions of patients that use biomedical ontologies to annotate phenotypes. Phenopackets are increasingly being used as a standard format for exchanging, aggregating and analysing human disease information [20]. We developed Klarigi to natively support the phenopackets format, converting it internally into the required data model, which we believe is a forward-looking facility for the anticipated future use of phenopackets, for example as an export format for EHRs.

To further evaluate Klarigi in the context of this data representation method, we examined a dataset of 384 rare disease patients whose phenotypes were originally reported in the literature, and have subsequently had their phenotypes transcribed into the phenopacket format [37]. The dataset describes a range of rare diseases, and in our use case, we examine the disease with the most patient descriptions, `Hypotonia, infantile, with psychomotor retardation and characteristic facies 3 (OMIM:616900)` (IHPRF3), with 19 patients. IHPRF3 is an ultra-rare genetic disorder broadly characterised by global developmental delay and regression, brain atrophy, extreme hypotonia and a wide range of characteristic facial dysmorphologies [41]. It is caused by recessive mutations in the TBCK gene [3] In the dataset we use, the patient descriptions are derived from four separate publications, listed in Table 7 were taken from these citations. The 2018-03-08 release of HPO was used to annotate these patients, and this was the version we used for the analysis.

**Table 7.** The publications that report the 19 patients with IHPRF3 that are phenotyped in the phenopackets dataset.

| Short Name | Full Title | Patients |
| --- | --- | --- |
| Bhoj 2016 [3] | Mutations in TBCK, Encoding TBC1-Domain-Containing Kinase, Lead to a Recognizable Syndrome of Intellectual Disability and Hypotonia | 10 |
| Chong 2016 [5] | Recessive Inactivating Mutations in TBCK, Encoding a Rab GTPase-Activating Protein, Cause Severe Infantile Syndromic Encephalopathy | 5 |
| Guerreiro 2016 [11] | Mutation of TBCK causes a rare recessive developmental disorder | 3 |
| Zapata-Aldana 2019 [50] | Further delineation of TBCK - Infantile hypotonia with psychomotor retardation and characteristic facies type 3 | 2 |
| Total | - | 19 |

Supplementary Table 5 shows the full set of 61 associated classes that constitute the univariate results. Meanwhile, Table 8 shows the multivariable results, whose overall inclusion describes complete coverage of the dataset with at least one of these classes, expressed in a much smaller explanatory set of 33 classes. Enrichment analysis was also performed using the Fisher and binomial tests, and the significant results of this analysis are shown in Supplementary Table 6.

**Table 8.** Multivariable results for IHPRF3 in Klarigi. Reclassification AUC was 1.

| OMIM:616900 (19 members) | r-score | Inclusion | Exclusion | IC |
| --- | --- | --- | --- | --- |
| Severe muscular hypotonia (HP:0006829) | 0.8 | 0.68 (p=0.001) | 0.95 (p=0.009) | 1.0 |
| Developmental regression (HP:0002376) | 0.63 | 0.58 (p=0.001) | 0.68 (p=0.007) | 1.0 |
| Severe global developmental delay (HP:0011344) | 0.5 | 0.42 (p=0.001) | 0.62 (p=0.012) | 1.0 |
| Abnormality of upper lip vermillion (HP:0011339) | 0.46 | 0.47 (p=0.001) | 0.45 (p=0.007) | 0.83 |
| Respiratory insufficiency (HP:0002093) | 0.42 | 0.42 (p=0.001) | 0.42 (p=0.042) | 0.81 |
| Reduced tendon reflexes (HP:0001315) | 0.42 | 0.53 (p=0.001) | 0.35 (p=0.018) | 0.82 |
| Small basal ganglia (HP:0012697) | 0.41 | 0.26 (p=0.001) | 0.95 (p=0.007) | 1.0 |
| Exaggerated cupid's bow (HP:0002263) | 0.41 | 0.26 (p=0.001) | 0.95 (p=0.009) | 1.0 |
| Macroglossia (HP:0000158) | 0.39 | 0.26 (p=0.001) | 0.78 (p=0.01) | 0.95 |
| Areflexia (HP:0001284) | 0.39 | 0.32 (p=0.001) | 0.5 (p=0.012) | 0.89 |
| Profound global developmental delay (HP:0012736) | 0.36 | 0.26 (p=0.001) | 0.58 (p=0.012) | 1.0 |
| Prominent nasal bridge (HP:0000426) | 0.36 | 0.26 (p=0.001) | 0.58 (p=0.014) | 1.0 |
| Skeletal muscle hypertrophy (HP:0003712) | 0.35 | 0.26 (p=0.001) | 0.51 (p=0.013) | 0.78 |
| Sloping forehead (HP:0000340) | 0.34 | 0.21 (p=0.001) | 0.95 (p=0.006) | 1.0 |
| Aplasia/Hypoplasia of the cerebellar vermis (HP:0006817) | 0.33 | 0.26 (p=0.001) | 0.45 (p=0.044) | 0.87 |
| Highly arched eyebrow (HP:0002553) | 0.33 | 0.26 (p=0.001) | 0.45 (p=0.039) | 1.0 |
| Cerebellar vermis hypoplasia (HP:0001320) | 0.33 | 0.26 (p=0.001) | 0.45 (p=0.044) | 0.95 |
| Tented upper lip vermillion (HP:0010804) | 0.31 | 0.21 (p=0.001) | 0.62 (p=0.015) | 1.0 |
| Coarse facial features (HP:0000280) | 0.29 | 0.26 (p=0.001) | 0.34 (p=0.036) | 1.0 |
| Small forehead (HP:0000350) | 0.29 | 0.26 (p=0.001) | 0.34 (p=0.041) | 1.0 |
| Narrow forehead (HP:0000341) | 0.29 | 0.26 (p=0.001) | 0.34 (p=0.041) | 1.0 |
| Aplasia/Hypoplasia of the corpus callosum (HP:0007370) | 0.29 | 0.37 (p=0.001) | 0.24 (p=0.094) | 0.95 |
| Aplasia/Hypoplasia of the cerebral white matter (HP:0012429) | 0.29 | 0.37 (p=0.001) | 0.24 (p=0.094) | 0.95 |
| Hypoplasia of the corpus callosum (HP:0002079) | 0.29 | 0.32 (p=0.001) | 0.27 (p=0.094) | 1.0 |
| Cerebral white matter hypoplasia (HP:0012430) | 0.29 | 0.32 (p=0.001) | 0.27 (p=0.094) | 1.0 |
| Brachycephaly (HP:0000248) | 0.29 | 0.21 (p=0.001) | 0.45 (p=0.036) | 0.95 |
| Absent speech (HP:0001344) | 0.28 | 0.37 (p=0.001) | 0.23 (p=0.085) | 1.0 |
| Abnormality of the cerebellar vermis (HP:0002334) | 0.27 | 0.26 (p=0.001) | 0.28 (p=0.042) | 0.8 |
| Diffuse cerebellar atrophy (HP:0100275) | 0.27 | 0.16 (p=0.001) | 0.95 (p=0.006) | 1.0 |
| Global brain atrophy (HP:0002283) | 0.27 | 0.16 (p=0.001) | 0.95 (p=0.006) | 1.0 |
| Global developmental delay (HP:0001263) | 0.26 | 0.95 (p=0.001) | 0.15 (p=0.055) | 0.89 |
| Open mouth (HP:0000194) | 0.26 | 0.16 (p=0.001) | 0.7 (p=0.011) | 1.0 |
| Diffuse cerebral atrophy (HP:0002506) | 0.26 | 0.16 (p=0.008) | 0.7 (p=0.002) | 1.0 |
| Visual impairment (HP:0000505) | 0.25 | 0.21 (p=0.001) | 0.31 (p=0.052) | 0.87 |
| *Overall* | - | *100.0* | - | - |

To provide a quantitative evaluation for this analysis involving a small set of patients, we compared them with the disease annotations defined by the HPO database. HPO database annotations were developed through a combination of expert curation and text mining from literature, clinical descriptions, and experimental evidence [28]. There are 46 such annotations, listed in Supplementary Table 4. To measure how characteristic the result sets are we calculated semantic similarity between each one and the set of HPO database annotations. We used the Resnik method of pairwise similarity, with the Zhou method of information content, with the best match average groupwise strategy. The results of the Fisher and Binomial tests are here treated separately, since neither forms a subset of the other. Table 9 lists the semantic similarity value for each comparison.

**Table 9.** Resnik+BMA+Zhou semantic similarity between the the explanatory set for each method and the definitive set of IHPRF3 HPO annotations defined by the HPO annotations database.

| Method | Similarity |
| --- | --- |
| Fisher | 0.560 |
| Binomial | 0.607 |
| Klarigi (u) | 0.703 |
| Klarigi (mv) | **0.72** |

We can also use Klarigi to evaluate alternative ways of grouping the data, or subsets of the data. In this use case, we explored 19 IHPRF3 patient data across different publications, shown in Table 7. In Supplementary Table 7 we show multivariable Klarigi results for IHPRF3 patients grouped by the publication in which they are reported.

## Discussion

We have described the design and implementation of Klarigi, and its application to two use-cases, one to the comparison of two phenotypically similar diseases from text-derived phenotypes, and another to rare disease patients from expert annotations created from literature. In this section, we will interpret and discuss the results, and provide more discussion of Klarigi in context, including limitations and future work.

### Interpretation of Results

#### Pulmonary Embolism

The large values of *overall_inclusion* calculated for the compositional results in Table 4 indicate a high overall coverage of the groups by the explanatory modules. There are many shared explanatory classes between the two diseases here, expressing their known similarity in presentation, as evidenced by previous compositional analysis of known presenting phenotypes for both conditions [32]. Half of the explanatory terms listed for pneumonia are shared by pulmonary embolism, often with very similar scores for *inclusion,* such as a difference of 0.05 for `dyspnea (HP:0002094)`. Differences in terms for this set may provide candidates for discriminatory terms, such as `cough (HP:0012735)` and `arryhthmia (HP:0011675)`. Indeed, the enrichment results include `cough (HP:0012735)` as a significantly over-represented class, being the only significant result produced by the binomial method.

The discriminatory analysis aims to produce explanatory sets that can be used to distinguish between the other groups in the dataset, shown in Table 5. We can see that none of the four redundant terms in the compositional results appear in either set of results, though non-redundant and highly compositional classes such as `cough (HP:0012735)` are preserved.

We see in the discriminatory analysis inclusion of classes describing known predisposing conditions for each disease. For example, pneumonia is a common complication of chronic renal disease [46] and PE is often a sequel to deep vein thrombosis [25] which is associated with lower limb pain, and obesity is similarly a known associated factor [49]. Tachycardia, hypercoagulability, and syncope are all established characteristics of PE, and it is notable that for pneumonia the most discriminating features are associated with respiratory difficulty; cough, wheezing, fever, and atrial arrhythmias. While fever may be found in some cases of PE it is clear that in our cohort this was not sufficiently common to contribute to the compositional analysis or to prevent it appearing as a discriminating class for pneumonia. The phenotype profiles derived from discharge letters do not contain information on D dimer levels [23], hence this frequently used measure did not appear in the results.

The discriminatory power of the sets is further explicated by the classification analysis, the results of which are given in Table 6. Reduction in performance for prediction of pneumonia with the Klarigi classifier was minimal, indicating that the set of explanatory terms generalised well to the phenotypic distribution of those patients in the test set. This reduction was greater for pulmonary embolism, which tracks the lower overall quality of this explanatory set evident on the training set, however the classifier still achieved a high performance.

For all classification methods and for both groups, models derived from the sets of explanatory terms generated by Klarigi outperformed those from enrichment. This is very strking in the case of pulmonary embolism, where performance of the enrichment on the test set approached random by all classification methods, with poor generalisation also evidenced by the very low overall inclusion value on the holdout set.

Another insight from the classification analysis on the test set is that the Klarigi classifier consistently performed substantially better than both semantic similarity approaches. This helps to provide evidence that the exclusion score, which makes up the Klarigi classification model, is a good measure of term discrimination, that generalises to use for quantifying discriminatory sets.

#### Rare Disease Annotation with Phenopackets

The phenopackets dataset is much smaller than the MIMIC dataset, describing far fewer patients. The annotations of patients were also derived by an expert from literature reports, instead of being mined from text. The task also differs, in that we are evaluating IHPRF3 patients against a large background distribution of patients with other rare genetic diseases, instead of comparing their phenotype profiles head-to-head with those of a single chosen, and phenotypically similar disease. This difference is evidenced by the increased clarity of the results, with large individual values for inclusion and exclusion, overall inclusion, and reclassification AUC. Klarigi was also able to identify a much smaller subset of the univariate sets during the multivariate stage, and this provides an example of the identification of sparse representations of entity groups, in the case that the representation of terms across the group is sufficiently homogenous.

The small number of disease cases also informed the decision not to perform a train-test validation for this use case. Instead, explanatory sets were compared with semantic similarity to the HPO database annotations for the disease. In this evaluation, Klarigi’s result sets were shown to be more similar to the definitional annotations than were both enrichment approaches.

We can also use this dataset to identify relationships that are not represented in the structured description of the disease. Several classes identified by the multivariate results, described in Table 8, are not found in the HPO database annotations. For instance, `brachycephaly (HP:0000248)` is not included, though it affects 21% of described patients with IHPRF3. As such, the findings produced by Klarigi could constitute new disease associations for the disease, not currently expressed in existing scientific databases describing it. In addition, of the HPO database annotations that are shared by Klarigi’s representation, we provide additional information about patient representation, information content, and how exclusive these phenotypes are for patients across a wide set of rare diseases, all information that can be used to enrich secondary use of these phenotype associations.

Supplementary Table 8 shows the multivariable Klarigi results for IHPRF3 when stratified by the publication they appear in. This analysis allows us to identify features that may be more specific to particular publications and the patient groups they report. For example, `intellectual disability (HP:0001249)` is over-represented in the ten patients described by Bhoj et al. [3], and both listed phenotypes for the three siblings in Guerreiro et al. [11] are highly exclusive and inclusive. The terms appearing in these sets are highly discriminatory, and reflect the genetic and phenotypic heterogeneity of the patient groups described. Interestingly, we can see from the metadata associated with the HPO database annotations, that some associations were directly evidenced by the OMIM clinical description, which in turn makes use of publications not represented in the phenopackets dataset, in particular Alazami et al. [1].

### Comparison to Enrichment Analysis

The results of Klarigi cannot easily be directly compared to enrichment analysis. Fundamentally, they solve different problems, since enrichment analysis methods only attempt to identify significantly associated classes, and usually only for univariate relationships. Meanwhile, Klarigi produces its results through class scoring and discrimination rather than a significance cut-off. Therefore, while our quantitative evaluations show that Klarigi out-performs enrichment in all cases, this primarily indicates the different problem that Klarigi solves: semantic explanation, or identification of characteristic and discriminatory sets of terms for an annotated entity group.

Our results can also be used to illustrate where enrichment does and does not perform well for semantic explanation. In the case of pneumonia, there are many individual classes with a strong and significant univariate association with the group, and this leads to a sizeable set of associated classes, which do function together as a discriminatory module, evident by the high performance on the test set, listed in Table 6. In the case of pulmonary embolism, however, the phenotypic distribution is more complex, and only a single term is identified. By contrast, Klarigi identifies a set of terms that, in combination, characterise the group. Here the individual significance of terms becomes less relevant, since they can nevertheless appear in a discriminative module, whose generalisability be checked through means other than a statistical test, such as through classification or measurement of *overall_inclusion.*

Not relying on significance testing for term identification also allows Klarigi to perform. The proof that these terms ‘fit’ the dataset can be. Furthermore, Klarigi provides individual significance testing, however even where exclusion scores for particular classes may not be significant, it is highly likely that the combination of explanatory classes will be. Since we are not selecting multivariable terms based on significance testing, this gives us the opportunity to manually and reactively modify hyper-parameters to identify better explanatory modules for the group. For example, if initial results are too general, or if the result set is dominated by a single class, the *max-inclusion* parameter can be modified.

Klarigi’s univariate mode is more comparable to enrichment, in that it identifies a set of pairwise relationships between terms and the group of interest. However, Klarigi provides more information about those relationships, and can therefore be used to more clearly explicate and interpret the dataset. For example, enrichment identified `congestive heart failure (HP:0002098)` as a highly over-represented class for pneumonia, with large values for odds ratio and z-score. This association is not incorrect, and in fact it is a strong discriminator, which also appears in the Klarigi solution. However, while its discriminatory power is high, it is not a very highly compositional term, with only 0.3 for *inclusion.* In the case of another highly associated class for pneumonia, `cough (HP:0012735)` appears to strongly characterise pneumonia, however this phenotype also appears with high prevalence in the pulmonary embolism group too, evidenced by the Klarigi-derived *inclusion* score of 0.25, compared with 0.67 for pneumonia. In cases like this, large values for zscore and odds ratio may be confusing or difficult to interpret, and may not provide enough information to those attempting to gain a holistic understanding of each group and their relationship with each other. Meanwhile, Klarigi captures this combination of facts in the scores it presents for `congestive heart failure (HP:0002098)` and `cough (HP:0012735)`.

Another distinguishing feature is Klarigi’s use of an ontology reasoner. Most enrichment methods do not make use of a reasoner, working instead on purely asserted axioms, or in some cases though pre-inferred knowledge graphs produced by preprocessing tools such as OntoFunc. Use of an ontology reasoner provides access to inferential reasoning and relationships between classes when identifying explanatory terms. For example, an additional inferred subclass relationship would affect *inclusion*, *exclusion*, and *specificity* scores as instances are determined across the ontology structure. Furthermore, use of an ontology reasoner confirms internal consistency of the ontology being used to perform the investigation.

### Context, Limitations, and Future Work

The design of Klarigi is a development upon principles first explored in our previous work, in which we developed an algorithm for deriving multivariable explanations for semantic clusters identified from Human Phenotype Ontology (HPO) phenotype profiles [40]. In this work, the approach is heavily modified and improved upon, including changes to the algorithm and scoring system, and is generalised to be applicable to any dataset and ontology. In particular, in the Design and Implementation section, we discussed limitations to the previous definition of the *exclusion* score, and created a new definition that is less sensitive to class imbalance.

While one feature of Klarigi is the ability to interactively modify hyperparameters to identify desirable result sets, these parameters also introduce complexity to the process, and understanding the interplay of the parameters and their effect on the results may require manual work, since it will differ depending on the dataset. Moreover, it may not be easily possible for a human operator to find the optimal parameter configuration.

Grid search could also be employed to automatically identify good settings for parameters, although a challenge here lies in identifying a good optimisation heuristic, as the current `overall_inclusion` may be equivalent across result sets of different quality, with respect to a desired outcome for the human operator, depending on their research goals - for example, five terms that describe 100% of the group may be preferable to one term that also describes 100% of the group. Such a search through possible overall parameters could also add significantly to the execution time. However, overcoming this problem would also help to avoid the problem of providing parameter defaults that perform well across different ontologies, datasets, and algorithmic parameters, and reduce the amount of human input required to obtain quality results. Parameter optimisation could also potentially introduce problems with multiple testing. We controlled for this, however, by choosing not to employ permutation testing until the final set of desired results is received - Klarigi currently requires an additional parameter to be passed for significance testing to be performed.

It is also possible that different scores could be introduced, or different definitions of existing scores could be explored. For example, the option to re-weight the r-score, in a similar manner to differential weight applications to precision and recall in an f-score.

The information Klarigi produces about unique characterisation of rare diseases may also benefit development of disease-phenotype associations, which are used in a wide range of analysis approaches, such as rare disease variant prioritisation or diagnosis. This contribution could consist simply in the identification of additional phenotypes through analysis of either larger aggregated datasets similar to the phenopackets dataset, or text-mined data from clinical encounters or literature. Moreover, we showed in Supplementary Table 8 that we were able to identify disease phenotypes that were more exclusively associated with single publications. Taking note of this information could support development of high quality disease-phenotype relationships. For example, particular phenotypes that are very exclusive to one or few publications may be less preferred as candidates for associations, or associations could be provided with a measure of inter-literature agreement. Such a dataset could be produced using a larger set of phenopackets data, or through mining literature and clinical narrative for phenotype profiles. Klarigi parameters could be configured to identify the necessary level of association desired to constitute a disease-phenotype link.

The measures described in our experiment could also be combined with enrichment analysis approaches. Enrichment analysis use a measure of *enrichment score*, which could comprise any of the r-score, inclusion, or exclusion scores given herein, then using the usual method of testing for significant effects. Indeed, the use of significance testing upon the final set of multivariable results can be considered as a kind of enrichment analysis. Other potential expansions of functionality could consist in extending the tool to facilitate a wider range of information content measures, involving multi-faceted semantic similarity, or identification of sub-groups through clustering.

Admissions sampling for the MIMIC dataset did not consider whether multiple admissions were attributed to the same patient, and this could potentially be a small source of bias. However, we do not believe it affects the conclusions of this study, since it would not favour a particular algorithm. In the same way, different publications described by the phenopackets dataset could potentially describe the same patients, although there is no way to confirm this programmatically (although this could conceivably be encoded using the phenopackets format). Future work could involve fully evaluating the use of Klarigi in a clinical classification setting, either standalone or as a feature selection methodology.

We did not compare the approach with another multivariable enrichment method. This is because existing implementations for multivariable enrichment are either limited to the Gene Ontology, or require a complicated setup process. The possibility for Klarigi to be easily applied to any ontology and ontology-annotated dataset is a major benefit of the approach. We addressed this by making these methodological differences clear in our problem description, comparison of the results, and discussion.

We anticipate that Klarigi can be used to perform improved semantic descriptive analysis of biomedical entities. This is an area of research that is of recent interest, especially with advancements in structured representation of phenotype profiles through phenopackets, and the automated creation of phenotypic profiles through text mining. For example, one study performed a statistical analysis of text-mined phenotype profiles for 4,095 individuals with Down Syndrome [14], descriptively reporting on phenotypes and their frequency of appearance. This analysis, however, only used a measure of patient frequency to stratify phenotypes. The use of Klarigi in this case would have introduced measures of representation for considered phenotypes, for example overall coverage of unique individuals, and the utility to easily switch groupings to explore the different aspects of the data.

## Conclusions

Klarigi provides a new way to solve the task of semantic explanation, and thus to explore and interpret biomedical datasets linked to ontologies. We have demonstrated that it can be used to gain insight into relationships between biomedical entities with clinical relevance.

## Supporting information

Supplemental Materials referred to in manuscript

## Funding

GVG and LTS acknowledge support from the NIHR Birmingham ECMC, NIHR Birmingham SRMRC, Nanocommons H2020-EU (731032), MAESTRIA (Grant agreement ID 965286), HYPERMARKER (Grant agreement ID 101095480), and the MRC Heath Data Research UK (HDRUK/CFC/01), an initiative funded by UK Research and Innovation, Department of Health and Social Care (England) and the devolved administrations, and leading medical research charities. The views expressed in this publication are those of the authors and not necessarily those of the NHS, the National Institute for Health Research, the Medical Research Council or the Department of Health. RH, PNS and GVG were supported by funding from King Abdullah University of Science and Technology (KAUST) Office of Sponsored Research (OSR) under Award No. URF/1/3790-01-01. AK was additionally supported by the Medical Research Council (MR/S003991/1).PNS and GVG acknowledge the support of The Alan Turing Institute.

## Abbreviations

HPO: Human Phenotype Ontology
IHPRF3: Hypotonia infantile with psychomotor retardation and characteristic facies 3
OMIM: Online Mendelian Inheritance In Man
OWL: Web Ontology Language
GSEA: Gene Set Enrichment Analysis
EGL: Exclusive Group Loading
GO: Gene Ontology
AUC: Area Under receiver operating Characteristic
IC: Information Content
MIMIC-III: Medical Information Mart for Intensive Care III
ML: Machine Learning.

## Ethics approval and consent to participate

This work makes use of the MIMIC-III dataset, which was approved for construction, de-identification, and sharing by the BIDMC and MIT institutional review boards (IRBs). Further details on MIMIC-III ethics are available from its original publication (DOI:10.1038/sdata.2016.35). Work was undertaken in accordance with the MIMIC-III guidelines. As such, while the code used to process MIMIC data is provided in the GitHub repository, the data itself is not made available.

## Competing interests

John Williams is an employee of Eisai, Inc. Eisai, Inc had no role in funding or design of this study.

## Authors’ contributions

**LTS** conceived of the study, designed and implemented the software, and created the initial manuscript. **JAW** contributed to design of the software, experimental design, and the manuscript, particularly surrounding the statistical analysis and significance testing. **AK** contributed to the manuscript and experimental design. **SR, SCP** tested multiple prototypical versions of the software and provided feedback that contributed to the final design. **PNS** performed clinical and biological interpretation of results, contributed to the manuscript and experimental design. **HF and SB** contributed to project supervision and experimental design. **RH and GVG** supervised the project, contributed to experimental design, and contributed to the manuscript. All authors approved the manuscript for submission.

## Notes

https://github.com/reality/klarigi/

https://github.com/reality/klarigi_paper/

### Summary of Updates

Resampled to correct class removals. Added clinical and biological interpretation. Rewritten aspects of discussion, results, and intro to clarify and provide better description of the approach and compare to other methods.

