## Supplemental Materials referred to in manuscript for "Klarigi: Characteristic Explanations for Semantic Data"

### Supplementary Material

#### Design and Implementation

**Data:** nPerm = Number of monte carlo permutations
AE = All unique classes in O that appear in E
**Result:** testDist = Sorted list of calculated test statistics, length = nPerm + 1
pValue = Calculated empirical p-value
**for** *n in nPerm* **do**
    $\text{ET}_j = |E|$
    **for** *p in E* **do**
        $\text{ET}_{jp} = random\_sample(AE, length(E_p))$
    **end**
    $\text{testDist}_n = calculate\_inclusion\_and\_exclusion(ET)$
**end**
;
testDist = rank(testDist);
r = length(testStat ≥ testDist);
pValue = $\frac{r+1}{nPerm+1}$;
return pValue;
**Algorithm 1:** Algorithm for generating p-values for inclusion and exclusion scores through permutation.

#### Use Case: Pulmonary Embolism

Table 1: All univariate scores for pulmonary embolism, derived by Klarigi with default parameters, using Resnik IC.

| **Class** | **Power** | **Inclusivity** | **Exclusivity** | **Specificity** |
|---|---|---|---|---|
| Sinus tachycardia (HP:0011703) | 0.17 | 0.25 | 0.13 | 1.0 |
| Increased body weight (HP:0004324) | 0.17 | 0.16 | 0.18 | 0.84 |
| Obesity (HP:0001513) | 0.16 | 0.14 | 0.17 | 0.87 |
| Abnormality of coagulation (HP:0001928) | 0.15 | 0.1 | 0.35 | 0.61 |
| Lower limb pain (HP:0012514) | 0.15 | 0.1 | 0.28 | 0.89 |
| Abnormal electrophysiology of sinoatrial node origin (HP:0011702) | 0.15 | 0.26 | 0.11 | 0.89 |
| Abnormality of musculoskeletal physiology (HP:0011843) | 0.15 | 0.14 | 0.17 | 0.45 |
| Limb pain (HP:0009763) | 0.15 | 0.1 | 0.27 | 0.84 |
| Pleuritic chest pain (HP:0033771) | 0.15 | 0.12 | 0.19 | 1.0 |
| Growth abnormality (HP:0001507) | 0.14 | 0.21 | 0.11 | 0.59 |
| Abnormality of body weight (HP:0004323) | 0.14 | 0.21 | 0.11 | 0.73 |
| Palpitations (HP:0001962) | 0.14 | 0.1 | 0.22 | 1.0 |
| Vascular dilatation (HP:0002617) | 0.13 | 0.1 | 0.19 | 0.74 |
| Syncope (HP:0001279) | 0.12 | 0.07 | 0.28 | 0.89 |
| Neoplasm by histology (HP:0011792) | 0.11 | 0.09 | 0.13 | 0.54 |
| Tachycardia (HP:0001649) | 0.11 | 0.29 | 0.06 | 0.74 |
| Arthritis (HP:0001369) | 0.11 | 0.22 | 0.07 | 0.74 |
| Hypercoagulability (HP:0100724) | 0.1 | 0.06 | 0.71 | 1.0 |
| Abnormal exteroceptive sensation (HP:0033747) | 0.1 | 0.06 | 0.29 | 0.8 |
| Somatic sensory dysfunction (HP:0003474) | 0.1 | 0.06 | 0.29 | 0.75 |

Table 2: All univariate scores for pneumonia, derived by Klarigi with default parameters, using Resnik IC.

| **Class** | **Power** | **Inclusivity** | **Exclusivity** | **Specificity** |
|---|---|---|---|---|
| Cough (HP:0012735) | 0.25 | 0.67 | 0.16 | 0.87 |
| Congestive heart failure (HP:0001635) | 0.22 | 0.3 | 0.17 | 0.95 |
| Respiratory distress (HP:0002098) | 0.21 | 0.34 | 0.15 | 0.92 |
| Respiratory insufficiency (HP:0002093) | 0.2 | 0.25 | 0.16 | 0.81 |
| Airway obstruction (HP:0006536) | 0.2 | 0.31 | 0.14 | 0.87 |
| Productive cough (HP:0031245) | 0.19 | 0.16 | 0.25 | 1.0 |
| Chronic pulmonary obstruction (HP:0006510) | 0.19 | 0.31 | 0.14 | 1.0 |
| Renal insufficiency (HP:0000083) | 0.19 | 0.37 | 0.13 | 0.84 |
| Respiratory failure (HP:0002878) | 0.19 | 0.22 | 0.16 | 1.0 |
| Chills (HP:0025143) | 0.19 | 0.27 | 0.14 | 1.0 |
| Wheezing (HP:0030828) | 0.19 | 0.38 | 0.12 | 1.0 |
| Pulmonary edema (HP:0100598) | 0.18 | 0.22 | 0.16 | 1.0 |
| Aspiration (HP:0002835) | 0.18 | 0.19 | 0.17 | 1.0 |
| Fever (HP:0001945) | 0.18 | 0.48 | 0.11 | 0.83 |
| Atrial fibrillation (HP:0005110) | 0.17 | 0.27 | 0.13 | 0.95 |
| Abnormality of temperature regulation (HP:0004370) | 0.17 | 0.48 | 0.1 | 0.77 |
| Confusion (HP:0001289) | 0.16 | 0.17 | 0.16 | 1.0 |
| Rhonchi (HP:0030831) | 0.16 | 0.2 | 0.14 | 1.0 |
| Atrial arrhythmia (HP:0001692) | 0.16 | 0.28 | 0.11 | 0.83 |
| Chronic kidney disease (HP:0012622) | 0.16 | 0.15 | 0.17 | 0.87 |
| Abnormal leukocyte count (HP:0011893) | 0.16 | 0.27 | 0.11 | 0.57 |
| Abnormal cellular immune system morphology (HP:0010987) | 0.15 | 0.29 | 0.1 | 0.51 |
| Abnormal leukocyte morphology (HP:0001881) | 0.15 | 0.29 | 0.1 | 0.51 |
| Stage 5 chronic kidney disease (HP:0003774) | 0.15 | 0.11 | 0.22 | 1.0 |
| Reduced consciousness/confusion (HP:0004372) | 0.15 | 0.24 | 0.11 | 0.8 |
| Abnormal renal physiology (HP:0012211) | 0.15 | 0.4 | 0.09 | 0.61 |
| Leukocytosis (HP:0001974) | 0.14 | 0.2 | 0.11 | 0.89 |
| Abnormal breath sound (HP:0030829) | 0.14 | 0.69 | 0.08 | 0.8 |
| Nonproductive cough (HP:0031246) | 0.14 | 0.1 | 0.24 | 1.0 |
| Supraventricular arrhythmia (HP:0005115) | 0.14 | 0.29 | 0.09 | 0.72 |
| Acute kidney injury (HP:0001919) | 0.14 | 0.18 | 0.11 | 1.0 |
| Abnormal lung morphology (HP:0002088) | 0.14 | 0.64 | 0.08 | 0.48 |
| Abnormal tracheobronchial morphology (HP:0005607) | 0.13 | 0.1 | 0.22 | 0.67 |
| Abnormal inflammatory response (HP:0012647) | 0.13 | 0.42 | 0.08 | 0.51 |
| Increased inflammatory response (HP:0012649) | 0.13 | 0.42 | 0.08 | 0.51 |
| Abnormal bronchus morphology (HP:0025426) | 0.13 | 0.09 | 0.22 | 0.72 |
| Crackles (HP:0030830) | 0.13 | 0.42 | 0.08 | 0.92 |
| Abnormal immune system morphology (HP:0032251) | 0.13 | 0.32 | 0.08 | 0.5 |
| Delirium (HP:0031258) | 0.12 | 0.09 | 0.2 | 1.0 |
| Sepsis (HP:0100806) | 0.12 | 0.13 | 0.12 | 1.0 |
| Tachypnea (HP:0002789) | 0.12 | 0.12 | 0.12 | 1.0 |
| Emphysema (HP:0002097) | 0.12 | 0.11 | 0.13 | 0.85 |
| Fatigue (HP:0012378) | 0.12 | 0.18 | 0.09 | 0.89 |
| Abnormality of the urinary system physiology (HP:0011277) | 0.11 | 0.47 | 0.07 | 0.41 |
| Abnormal heart valve morphology (HP:0001654) | 0.11 | 0.09 | 0.17 | 0.64 |
| Abnormal pulmonary interstitial morphology (HP:0006530) | 0.11 | 0.07 | 0.24 | 0.65 |
| Abnormality of thyroid physiology (HP:0002926) | 0.11 | 0.16 | 0.08 | 0.69 |
| Myocardial infarction (HP:0001658) | 0.11 | 0.13 | 0.09 | 1.0 |
| Abnormality of the upper respiratory tract (HP:0002087) | 0.11 | 0.09 | 0.14 | 0.61 |

**Table 2 – continued from previous page**

| **Class** | **Power** | **Inclusivity** | **Exclusivity** | **Specificity** |
|---|---|---|---|---|
| Malaise (HP:0033834) | 0.11 | 0.07 | 0.22 | 1.0 |
| Abnormality of the thyroid gland (HP:0000820) | 0.11 | 0.17 | 0.08 | 0.62 |
| Abnormal central motor function (HP:0011442) | 0.11 | 0.13 | 0.09 | 0.52 |
| Heart block (HP:0012722) | 0.11 | 0.08 | 0.14 | 0.78 |
| Cardiac conduction abnormality (HP:0031546) | 0.11 | 0.08 | 0.14 | 0.77 |
| Abnormal aortic valve morphology (HP:0001646) | 0.1 | 0.07 | 0.24 | 0.77 |
| Aortic valve stenosis (HP:0001650) | 0.1 | 0.07 | 0.24 | 1.0 |
| Hypothyroidism (HP:0000821) | 0.1 | 0.15 | 0.08 | 0.87 |
| Acute coronary syndrome (HP:0033678) | 0.1 | 0.13 | 0.09 | 1.0 |
| Insomnia (HP:0100785) | 0.1 | 0.11 | 0.1 | 0.92 |
| Hypoxemia (HP:0012418) | 0.1 | 0.08 | 0.14 | 0.95 |

Table 3: Enrichment results for pneumonia and pulmonary embolism

| | Binomial | | | Fisher | | |
|---|---|---|---|---|---|---|
| | zscore | OR | p | zscore | OR | p |
| Abnormality of coagulation (HP:0001928) | | | | 4.71 | 4.68 | 0.0051 |

| | Binomial | | | Fisher | | |
|---|---|---|---|---|---|---|
| | zscore | OR | p | zscore | OR | p |
| Cough (HP:0012735) | 10.2 | 5.66 | 1.3e-05 | 10.2 | 5.66 | 6.7e-22 |
| Congestive heart failure (HP:0001635) | | | | 5.93 | 3.82 | 1.8e-07 |
| Respiratory distress (HP:0002098) | | | | 5.72 | 0.403 | 1.6e-06 |
| Abnormal breath sound (HP:0030829) | | | | 5.45 | 2.44 | 3.4e-05 |
| Airway obstruction (HP:0006536) | | | | 4.85 | 2.75 | 0.00022 |
| Abnormal respiratory system morphology (HP:0012252) | | | | 4.83 | 1.04 | 0.001 |
| Chills (HP:0025143) | | | | 4.81 | 2.9 | 0.00024 |
| Respiratory insufficiency (HP:0002093) | | | | 4.74 | 3.12 | 0.00028 |
| Abnormality of temperature regulation (HP:0004370) | | | | 4.54 | 0 | 0.0022 |
| Aspiration (HP:0002835) | | | | 4.08 | 3.1 | 0.0074 |
| Pulmonary edema (HP:0100598) | | | | 3.99 | 2.79 | 0.014 |

#### Use Case: Phenopackets

Table 4: All univariate scores for IHPRF3 (OMIM:616900).

| **Class** | **r-score** | **Inclusivity** | **Exclusivity** | **IC** |
|---|---|---|---|---|
| Severe muscular hypotonia (HP:0006829) | 0.8 | 0.68 | 0.95 | 1.0 |
| Developmental regression (HP:0002376) | 0.63 | 0.58 | 0.68 | 1.0 |
| Severe global developmental delay (HP:0011344) | 0.5 | 0.42 | 0.62 | 1.0 |
| Abnormality of upper lip vermillion (HP:0011339) | 0.46 | 0.47 | 0.45 | 0.83 |
| Respiratory insufficiency (HP:0002093) | 0.42 | 0.42 | 0.42 | 0.81 |
| Reduced tendon reflexes (HP:0001315) | 0.42 | 0.53 | 0.35 | 0.82 |
| Small basal ganglia (HP:0012697) | 0.41 | 0.26 | 0.95 | 1.0 |
| Exaggerated cupid's bow (HP:0002263) | 0.41 | 0.26 | 0.95 | 1.0 |
| Macroglossia (HP:0000158) | 0.39 | 0.26 | 0.78 | 0.95 |
| Areflexia (HP:0001284) | 0.39 | 0.32 | 0.5 | 0.89 |
| Abnormality of the basal ganglia (HP:0002134) | 0.38 | 0.26 | 0.66 | 0.69 |
| Profound global developmental delay (HP:0012736) | 0.36 | 0.26 | 0.58 | 1.0 |
| Prominent nasal bridge (HP:0000426) | 0.36 | 0.26 | 0.58 | 1.0 |
| Skeletal muscle hypertrophy (HP:0003712) | 0.35 | 0.26 | 0.51 | 0.78 |
| Sloping forehead (HP:0000340) | 0.34 | 0.21 | 0.95 | 1.0 |

**Table 4 – continued from previous page**

| Class | r-score | Inclusivity | Exclusivity | IC |
|---|---|---|---|---|
| Aplasia/Hypoplasia of the cerebellar vermis (HP:0006817) | 0.33 | 0.26 | 0.45 | 0.87 |
| Highly arched eyebrow (HP:0002553) | 0.33 | 0.26 | 0.45 | 1.0 |
| Cerebellar vermis hypoplasia (HP:0001320) | 0.33 | 0.26 | 0.45 | 0.95 |
| Tented upper lip vermilion (HP:0010804) | 0.31 | 0.21 | 0.62 | 1.0 |
| Coarse facial features (HP:0000280) | 0.29 | 0.26 | 0.34 | 1.0 |
| Small forehead (HP:0000350) | 0.29 | 0.26 | 0.34 | 1.0 |
| Narrow forehead (HP:0000341) | 0.29 | 0.26 | 0.34 | 1.0 |
| Aplasia/Hypoplasia of the corpus callosum (HP:0007370) | 0.29 | 0.37 | 0.24 | 0.95 |
| Aplasia/Hypoplasia of the cerebral white matter (HP:0012429) | 0.29 | 0.37 | 0.24 | 0.95 |
| EEG abnormality (HP:0002353) | 0.29 | 0.32 | 0.27 | 0.58 |
| Hypoplasia of the corpus callosum (HP:0002079) | 0.29 | 0.32 | 0.27 | 1.0 |
| Abnormality of muscle size (HP:0030236) | 0.29 | 0.32 | 0.27 | 0.63 |
| Cerebral white matter hypoplasia (HP:0012430) | 0.29 | 0.32 | 0.27 | 1.0 |
| Brachycephaly (HP:0000248) | 0.29 | 0.21 | 0.45 | 0.95 |
| Absent speech (HP:0001344) | 0.28 | 0.37 | 0.23 | 1.0 |
| Abnormality of the cerebellar vermis (HP:0002334) | 0.27 | 0.26 | 0.28 | 0.8 |
| Diffuse cerebellar atrophy (HP:0100275) | 0.27 | 0.16 | 0.95 | 1.0 |
| Global brain atrophy (HP:0002283) | 0.27 | 0.16 | 0.95 | 1.0 |
| Abnormality of central nervous system electrophysiology (HP:0030178) | 0.27 | 0.32 | 0.24 | 0.56 |
| Global developmental delay (HP:0001263) | 0.26 | 0.95 | 0.15 | 0.89 |
| Open mouth (HP:0000194) | 0.26 | 0.16 | 0.7 | 1.0 |
| Diffuse cerebral atrophy (HP:0002506) | 0.26 | 0.16 | 0.7 | 1.0 |
| Seizures (HP:0001250) | 0.26 | 0.79 | 0.15 | 0.62 |
| Visual impairment (HP:0000505) | 0.25 | 0.21 | 0.31 | 0.87 |
| Shallow orbits (HP:0000586) | 0.25 | 0.16 | 0.55 | 1.0 |
| Abnormality of bony orbit of skull (HP:3000030) | 0.25 | 0.16 | 0.55 | 0.95 |
| Partial agenesis of the corpus callosum (HP:0001338) | 0.25 | 0.16 | 0.55 | 1.0 |
| Abnormality of the corpus callosum (HP:0001273) | 0.25 | 0.37 | 0.18 | 0.83 |
| Abnormality of upper lip (HP:0000177) | 0.24 | 0.47 | 0.16 | 0.69 |
| Cerebellar malformation (HP:0002438) | 0.24 | 0.26 | 0.23 | 0.74 |
| Muscular hypotonia (HP:0001252) | 0.24 | 0.95 | 0.14 | 0.8 |
| Abnormality of the periorbital region (HP:0000606) | 0.24 | 0.53 | 0.15 | 0.65 |
| Deeply set eye (HP:0000490) | 0.24 | 0.32 | 0.19 | 1.0 |
| Abnormal tongue morphology (HP:0030809) | 0.24 | 0.26 | 0.21 | 0.69 |
| Abnormality of the tongue (HP:0000157) | 0.24 | 0.26 | 0.21 | 0.65 |
| Cerebral atrophy (HP:0002059) | 0.23 | 0.21 | 0.26 | 0.82 |
| Abnormality of the lip (HP:0000159) | 0.23 | 0.53 | 0.15 | 0.62 |
| Focal seizures (HP:0007359) | 0.22 | 0.16 | 0.38 | 0.69 |
| Agenesis of corpus callosum (HP:0001274) | 0.22 | 0.16 | 0.38 | 1.0 |
| Osteoporosis (HP:0000939) | 0.22 | 0.16 | 0.38 | 0.92 |
| Atrophy/Degeneration affecting the cerebrum (HP:0007369) | 0.22 | 0.21 | 0.24 | 0.78 |
| Ventriculomegaly (HP:0002119) | 0.22 | 0.32 | 0.16 | 0.81 |
| Generalized seizures (HP:0002197) | 0.21 | 0.21 | 0.22 | 0.77 |
| Abnormality of the external nose (HP:0010938) | 0.21 | 0.42 | 0.14 | 0.67 |
| Abnormality of the cerebral white matter (HP:0002500) | 0.21 | 0.37 | 0.14 | 0.68 |
| Abnormal nervous system electrophysiology (HP:0001311) | 0.2 | 0.32 | 0.15 | 0.56 |
| Abnormality of the forehead (HP:0000290) | 0.2 | 0.53 | 0.13 | 0.7 |

Table 5: Enrichment results for IHPRF3 (OMIM:616900)

| | Binomial | | | Fisher | | |
|---|---|---|---|---|---|---|
| | zscore | OR | p | zscore | OR | p |
| Abnormality of upper lip vermillion (HP:0011339) | 9 | 4.14 | 4.7e-07 | 3.86 | Inf | 0.015 |
| Respiratory insufficiency (HP:0002093) | 8.17 | 1.99 | 5.8e-06 | 3.96 | 1.99 | 0.015 |
| Severe muscular hypotonia (HP:0006829) | 7.98 | Inf | 2e-07 | 7.98 | Inf | 3.3e-10 |
| Developmental regression (HP:0002376) | 7.76 | 0 | 1.5e-06 | 7.76 | 0 | 3.2e-07 |
| Muscular hypotonia (HP:0001252) | 7.23 | 0 | 4.4e-10 | | | |
| Cerebellar vermis hypoplasia (HP:0001320) | 6.64 | 24.9 | 0.00091 | | | |
| Aplasia/Hypoplasia of the cerebellar vermis (HP:0006817) | 6.64 | 24.9 | 0.00091 | | | |
| Reduced tendon reflexes (HP:0001315) | 6.23 | 14.7 | 8.2e-05 | 6.23 | 14.7 | 0.00017 |
| Abnormal muscle tone (HP:0003808) | 6.07 | 0 | 4.4e-08 | | | |
| EEG abnormality (HP:0002353) | 5.47 | 0 | 0.0031 | | | |
| Abnormality of the cerebellar vermis (HP:0002334) | 5.16 | 0.879 | 0.0089 | | | |
| Abnormality of muscle physiology (HP:0011804) | 5.14 | 0.172 | 2e-06 | | | |
| Abnormality of central nervous system electrophysiology (HP:0030178) | 5.12 | 0 | 0.0059 | | | |
| Visual impairment (HP:0000505) | 4.86 | 13.3 | 0.023 | | | |
| Abnormality of the corpus callosum (HP:0001273) | 4.82 | 0 | 0.0069 | | | |
| Seizures (HP:0001250) | 4.69 | 0 | 0.00037 | 4.69 | 0 | 0.0014 |
| Abnormality of the musculature (HP:0003011) | 4.5 | 24.7 | 3.2e-05 | 4.5 | 24.7 | 0.00046 |
| Abnormality of the tongue (HP:0000157) | 4.39 | 12.3 | 0.033 | | | |
| Abnormal tongue morphology (HP:0030809) | 4.39 | 0 | 0.033 | | | |
| Severe global developmental delay (HP:0011344) | 4.28 | 44.8 | 0.032 | 4.28 | 44.8 | 0.026 |
| Functional respiratory abnormality (HP:0002795) | 4.21 | 0 | 0.018 | | | |
| Abnormality of the cerebral white matter (HP:0002500) | 4.2 | 6.68 | 0.025 | | | |
| Ventriculomegaly (HP:0002119) | 4.16 | 0 | 0.036 | | | |
| Absent speech (HP:0001344) | 3.51 | 11.8 | 0.034 | | | |
| Abnormality of the head (HP:0000234) | 3.47 | Inf | 0.0024 | | | |
| Abnormality of muscle size (HP:0030236) | 3.45 | 0 | 0.026 | | | |
| Abnormality of head or neck (HP:0000152) | 3.39 | 1.87 | 0.0035 | 3.39 | 1.87 | 0.038 |
| Aplasia/Hypoplasia of the corpus callosum (HP:0007370) | 3.22 | 0.344 | 0.038 | | | |
| Neurodevelopmental abnormality (HP:0012759) | 3.11 | 0 | 0 | | | |
| Abnormality of the nervous system (HP:0000707) | 2.93 | 0 | 0 | | | |
| Neurodevelopmental delay (HP:0012758) | 2.29 | 0 | 0 | | | |
| Abnormality of nervous system physiology (HP:0012638) | 1.6 | Inf | 0 | | | |

Table 6: HPO database phenotype annotations for IHPRF3 (OMIM:616900)

| **Term ID** | **Label** |
|---|---|
| HP:0002465 | Poor speech |
| HP:0010945 | Fetal pyelectasis |
| HP:0001298 | Encephalopathy |
| HP:0001250 | Seizure |
| HP:0001265 | Hyporeflexia |
| HP:0001263 | Global developmental delay |
| HP:0002553 | Highly arched eyebrow |
| HP:0002518 | Abnormal periventricular white matter morphology |
| HP:0000007 | Autosomal recessive inheritance |
| HP:0001320 | Cerebellar vermis hypoplasia |
| HP:0002650 | Scoliosis |
| HP:0001321 | Cerebellar hypoplasia |
| HP:0000158 | Macroglossia |
| HP:0002093 | Respiratory insufficiency |
| HP:0002079 | Hypoplasia of the corpus callosum |
| HP:0002059 | Cerebral atrophy |
| HP:0002119 | Ventriculomegaly |
| HP:0002263 | Exaggerated cupid's bow |
| HP:0100704 | Cerebral visual impairment |
| HP:0002376 | Developmental regression |
| HP:0010804 | Tented upper lip vermilion |
| HP:0006829 | Severe muscular hypotonia |
| HP:0012697 | Small basal ganglia |
| HP:0006989 | Dysplastic corpus callosum |
| HP:0006970 | Periventricular leukomalacia |
| HP:0012736 | Profound global developmental delay |
| HP:0000750 | Delayed speech and language development |
| HP:0012708 | Reduced brain N-acetyl aspartate level by MRS |
| HP:0011471 | Gastrostomy tube feeding in infancy |
| HP:0000286 | Epicanthus |
| HP:0000280 | Coarse facial features |
| HP:0000256 | Macrocephaly |
| HP:0000212 | Gingival overgrowth |
| HP:0001562 | Oligohydramnios |
| HP:0001558 | Decreased fetal movement |
| HP:0001500 | Broad finger |
| HP:0000341 | Narrow forehead |
| HP:0000340 | Sloping forehead |
| HP:0012471 | Thick vermilion border |
| HP:0000490 | Deeply set eye |
| HP:0000463 | Anteverted nares |
| HP:0012444 | Brain atrophy |
| HP:0000414 | Bulbous nose |
| HP:0000426 | Prominent nasal bridge |
| HP:0001837 | Broad toe |
| HP:0012510 | Extra-axial cerebrospinal fluid accumulation |

Table 7: Multivariable Klarigi results for the 19 patients with IHPRF3 described in the phenopackets dataset, grouped by the publication they are described in.

| **Chong 2016 (5 members)** | **r-score** | **Inclusion** | **Exclusion** | **IC** |
|---|---|---|---|---|
| Profound global developmental delay (HP:0012736) | 0.85 | 1.0 | 0.74 | 1.0 |
| Prominent nasal bridge (HP:0000426) | 0.85 | 1.0 | 0.74 | 1.0 |
| Abnormality of the nasal alae (HP:0000429) | 0.85 | 1.0 | 0.74 | 0.82 |
| Abnormality of the nares (HP:0005288) | 0.85 | 1.0 | 0.74 | 0.82 |
| Feeding difficulties (HP:0011968) | 0.85 | 1.0 | 0.74 | 0.83 |
| Exaggerated cupid's bow (HP:0002263) | 0.85 | 1.0 | 0.74 | 1.0 |
| Cerebellar vermis hypoplasia (HP:0001320) | 0.85 | 1.0 | 0.74 | 0.95 |
| Small forehead (HP:0000350) | 0.85 | 1.0 | 0.74 | 1.0 |
| Small basal ganglia (HP:0012697) | 0.85 | 1.0 | 0.74 | 1.0 |
| Narrow forehead (HP:0000341) | 0.85 | 1.0 | 0.74 | 1.0 |
| Anteverted nares (HP:0000463) | 0.85 | 1.0 | 0.74 | 0.95 |
| Aplasia/Hypoplasia of the cerebellar vermis (HP:0006817) | 0.85 | 1.0 | 0.74 | 0.87 |
| Abnormality of the cerebellar vermis (HP:0002334) | 0.85 | 1.0 | 0.74 | 0.8 |
| Cerebellar malformation (HP:0002438) | 0.85 | 1.0 | 0.74 | 0.74 |
| Highly arched eyebrow (HP:0002553) | 0.85 | 1.0 | 0.74 | 1.0 |
| *Overall* | - | *100.0* | - | - |
| **Zapata-Aldana 2019 (2 members)** | **r-score** | **Inclusion** | **Exclusion** | **IC** |
| Impaired social interactions (HP:0000735) | 0.94 | 1.0 | 0.89 | 0.87 |
| Poor eye contact (HP:0000817) | 0.94 | 1.0 | 0.89 | 1.0 |
| Respiratory failure (HP:0002878) | 0.94 | 1.0 | 0.89 | 1.0 |
| Abnormal social behavior (HP:0012433) | 0.94 | 1.0 | 0.89 | 0.81 |
| Long philtrum (HP:0000343) | 0.94 | 1.0 | 0.89 | 1.0 |
| Abnormality of the philtrum (HP:0000288) | 0.94 | 1.0 | 0.89 | 0.79 |
| *Overall* | - | *100.0* | - | - |
| **Bhoj 2016 (10 members)** | **r-score** | **Inclusion** | **Exclusion** | **IC** |
| Cognitive impairment (HP:0100543) | 0.6 | 0.8 | 0.47 | 0.77 |
| Intellectual disability (HP:0001249) | 0.6 | 0.8 | 0.47 | 0.85 |
| Severe global developmental delay (HP:0011344) | 0.47 | 0.7 | 0.35 | 1.0 |
| *Overall* | - | *100.0* | - | - |
| **PMID:27275012-Guerreiro-2016-TBCK (2 members)** | **r-score** | **Inclusion** | **Exclusion** | **IC** |
| Early onset of sexual maturation (HP:0100000) | 0.94 | 1.0 | 0.89 | 0.83 |
| Overlapping toe (HP:0001845) | 0.94 | 1.0 | 0.89 | 1.0 |
| Deep palmar crease (HP:0006191) | 0.94 | 1.0 | 0.89 | 1.0 |
| Single transverse palmar crease (HP:0000954) | 0.94 | 1.0 | 0.89 | 1.0 |
| Abnormal dermatoglyphics (HP:0007477) | 0.94 | 1.0 | 0.89 | 0.72 |
| Precocious puberty (HP:0000826) | 0.94 | 1.0 | 0.89 | 0.85 |
| Infantile muscular hypotonia (HP:0008947) | 0.94 | 1.0 | 0.89 | 1.0 |
| Abnormality of the palmar creases (HP:0010490) | 0.94 | 1.0 | 0.89 | 0.82 |
| Abnormal palmar dermatoglyphics (HP:0001018) | 0.94 | 1.0 | 0.89 | 0.74 |
| *Overall* | - | *100.0* | - | - |
